# BacStalk: a comprehensive and interactive image analysis software tool for bacterial cell biology

**DOI:** 10.1101/360230

**Authors:** Raimo Hartmann, Muriel C.F. van Teeseling, Martin Thanbichler, Knut Drescher

**Author notes:** These authors contributed equally.

## Abstract

Prokaryotes display a remarkable spatiotemporal organization of processes within individual cells. Investigations of the underlying mechanisms rely extensively on the analysis of microscopy images. Advanced image analysis software has revolutionized the cell-biological studies of established model organisms with largely symmetric rod-like cell shapes. However’ algorithms suitable for analyzing features of morphologically more complex model species are lacking’ although such unusually shaped organisms have emerged as treasure-troves of new molecular mechanisms and diversity in prokaryotic cell biology. To address this problem’ we developed BacStalk’ a simple’ interactive’ and easy-to-use MatLab-based software tool for quantitatively analyzing images of commonly and uncommonly shaped bacteria’ including stalked (budding) bacteria. BacStalk automatically detects the separate parts of the cells (cell body’ stalk’ bud’ or appendage) as well as their connections’ thereby allowing in-depth analyses of the organization of morphologically complex bacteria over time. BacStalk features the generation and visualization of concatenated fluorescence profiles along cells’ stalks’ appendages’ and buds to trace the spatiotemporal dynamics of fluorescent markers. Cells are interactively linked to demographs’ kymographs’ cell lineage analyses’ and scatterplots’ which enables intuitive and fast data exploration and’ thus’ significantly speeds up the image analysis process. Furthermore’ BacStalk introduces a 2D representation of demo- and kymographs’ enabling data representations in which the two spatial dimensions of the cell are preserved. The software was developed to handle large data sets and to generate publication-grade figures that can be easily edited. BacStalk therefore provides an advanced image analysis platform that extends the spectrum of model organisms for prokaryotic cell biology to bacteria with multiple morphologies and life cycles.

**IMPORTANCE:** Prokaryotic cells show a striking degree of subcellular organization. Studies of the underlying mechanisms and their variation among different species greatly enhance our understanding of prokaryotic cell biology. The image analysis software tool BacStalk extracts an unprecedented amount of information from images of stalked bacteria, by generating interactive demographs, kymographs, cell lineages, and scatter plots that aid fast and thorough data analysis and representation. Notably, BacStalk can preserve the two spatial dimensions of cells when generating demographs and kymographs to accurately and intuitively reflect the intracellular organization. BacStalk also performs well on established, non-stalked model organisms with common or uncommon shapes. BacStalk therefore contributes to the advancement of prokaryotic cell biology, as it widens the spectrum of easily accessible model organisms and enables a more intuitive and interactive data analysis and visualization.

## INTRODUCTION

The study of prokaryotic model species has greatly contributed to the advancement of cell biology. Originally viewed as simple organisms lacking internal structure, prokaryotes are now acknowledged to display an impressive degree of subcellular organization despite their small size (1). The central question in cell biology is how cells spatiotemporally regulate the positioning of (macro)molecular machineries and how they control these complexes in order to grow, divide, and adequately respond to changes in their extracellular environment. Studies in established prokaryotic model organisms, such as *Bacillus subtilis*, *Escherichia coli*, *Vibrio cholerae*, and *Caulobacter crescentus*, have yielded many insights into the mechanisms that spatiotemporally organize prokaryotic cells. However, these species only represent a small part of the highly diverse prokaryotic world. Therefore, it is not surprising that studies of less commonly studied model species have recently uncovered a series of novel cell-biological pathways. These findings have significantly improved our understanding of prokaryotic cell biology and revealed that prokaryotes have often evolved multiple solutions to the same regulatory problem (2, 3).

A fundamental cell-biological process is cell division. The investigation of a range of different model species unveiled that this process is controlled by common, yet also diverse molecular mechanisms. In *E. coli* and *B. subtilis*, the proper division site placement is achieved through the Min (4) and nucleoid occlusion (NO) systems (5), which inhibit the assem bly of the cytokinetic FtsZ ring at inappropriate places (e.g. the cell poles) and times (e.g. before chromosome segregation has finished). *C. crescentus* also regulates Z-ring placement in a negative manner, but integrates the functions of both the Min and NO systems into a single protein, MipZ (6, 7). Studies of other organisms, such as *Streptomyces coelicolor* (8), *Streptococcus pneumoniae* (9, 10), and *Myxococcus xanthus* (11, 12) showed that the cell division machinery can also be positioned *via* positive regulation, i.e. through the active recruitment of FtsZ to the future cell division site. In the overwhelming majority of cases, cell division has been studied in species that divide through symmetric binary fission. However, there are various other ways by which prokaryotes grow and divide (13, 14), but little is known about the mechanisms steering and regulating these alternative cell division processes.

A nearly unexplored mode of cell division is stalk-terminal budding, in which cells produce one or multiple cytoplasm-filled extrusions of the cell envelope (termed prosthecae or stalks), the tips of which develop into daughter cells (15). Stalk-terminal budding occurs within several alphaproteobacterial species, and it appears to have arisen at least twice during evolution in the *Hyphomicrobium/Rhodomicrobium* and *Hyphomonas/Hirschia* lineages, respectively (16). Stalked bacteria provide a treasure trove for the analysis of cell division, as this group also includes organisms that bud directly from the mother cell or divide by binary fission (17–19). Moreover, many stalked bacteria undergo cell cycle-dependent changes in their morphology and physiology (20–25), and thereby provide critical new insights into the mechanisms that spatiotemporally organize bacterial cells (26). Indeed, one of the most prominent model organisms in the field of bacterial cell biology, *C. crescentus*, is also a stalked bacterium.

Primary characteristics of a suitable model organism for prokaryotic cell biology are: (i) ease of cultivation, (ii) availability of a genetic toolbox, and (iii) the ability to extract quantitative phenotypic information *via* light microscopy, aided by an efficient image analysis software. Although a variety of bacterial species are available that are easy to cultivate and genetically amenable, automated image analysis software is still limiting. The development of dedicated software tools, such as MicrobeTracker (27), Oufti (28), and MicrobeJ (29), and others (30–32), has enabled the efficient and automated analysis of light microscopy images of common prokaroytic model organisms to the point that even impressive high-throughput screening of mutant libraries based on morphology and intracellular organization are now possible (33). Although these software packages are becoming more versatile, they lack the possibility to extract quantitative information from prokaryotes with special morphologies. In the case of stalked bacteria, for instance, the detection of stalks in microscopy images routinely still proceeds manually. Existing image analysis algorithms usually fail to reliably recognize stalks because they are hardly visible and characterized by a very low contrast in phase contrast images, making automated segmentation challenging. When, as a work-around, stalks are fluorescently labeled to facilitate their visualization, their dimensions are typically increased significantly by diffraction. Although the latest version of MicrobeJ can relate filamentous structures to corresponding cells if they are fluorescently labelled and segmentable by thresholding (29), the possibilities for analyzing stalked bacteria are very limited. At present, it is for instance not possible to visualize concatenated fluorescence profiles along cells and stalks. In addition, the image analysis of stalked budding bacteria remains very difficult as the mother cell, stalk, and bud cannot be detected as connected entities that make up a single cell.

To provide a powerful image analysis software for prokaryotes with complex morphologies, such as stalked bacteria, we developed BacStalk: a ready-to-use MatLab-based software tool specialized for automated, label-free, time-resolved image analysis of stalked and non-stalked bacteria.

Using an intuitive graphical user interface (GUI), BacStalk enables the detection of fluorescence patterns and the morphometric analysis of cell bodies, and, if present, their corresponding stalks and buds. Importantly, BacStalk can visualize fluorescence signals in (populations of) cells in interactive demographs and kymographs in which the cells can be aligned according to relevant morphological features, and cell morphologies are visualized in addition to the fluorescence signals. This paper discusses the capabilities and functionality of BacStalk, based on image datasets of different bacterial species.

## RESULTS & DISCUSSION

BacStalk is designed for label-free bacterial cell and stalk detection in phase contrast images at pixel accuracy. As stalks are usually very dim and hardly visible, conventional thresholding approaches fail to separate stalks reliably from the image background. BacStalk overcomes these difficulties by implementing a two-step approach: First, cells are identified by enhancing image features of a typical length scale, followed by automatic thresholding and finding of connected components. In the second step, BacStalk detects connected stalks by performing local morphological operations. In detail, the neighboring pixels of each cell are scanned for subtle intensity minima indicating a possible stalk attachment point (Fig. 1A). If such a stalk anchor is found in terms of the lowest intensity value below a defined z-score (i.e. the number of standard deviations that a value deviates from the mean) along the expanded bacterial outline, the stalk backbone is generated by repeated dilation starting from the stalk attachment point while looking for other pixels in a defined range that are again below a defined z-score (Fig. 1A). If another cell is encountered during stalk propagation, both the initial cell and the touched counterpart are defined as related and can be treated as a connected structure (mother cell and bud) during cell tracking and the downstream analysis. The larger cell is hereby defined as mother cell. In cells and buds of stalked budding bacteria, the cell polarity is clearly defined by the stalk attachment point and indicated by a yellow dot at the end of the cell’s medial axis (Fig. 1A). In the case of stalk-free swarmer cells, the identity of the cell pole is guessed based on the cell morphology or the existence of intensity features, but can also be interactively changed manually at any stage of the analysis. As unambiguous stalk assignment in clumps of multiple cells is mostly impossible, cell clumps are automatically recognized and excluded from further analysis. The approach described above is relatively insensitive to uneven background illumination in the microscopy images and does not require any background correction.

**FIG 1.**
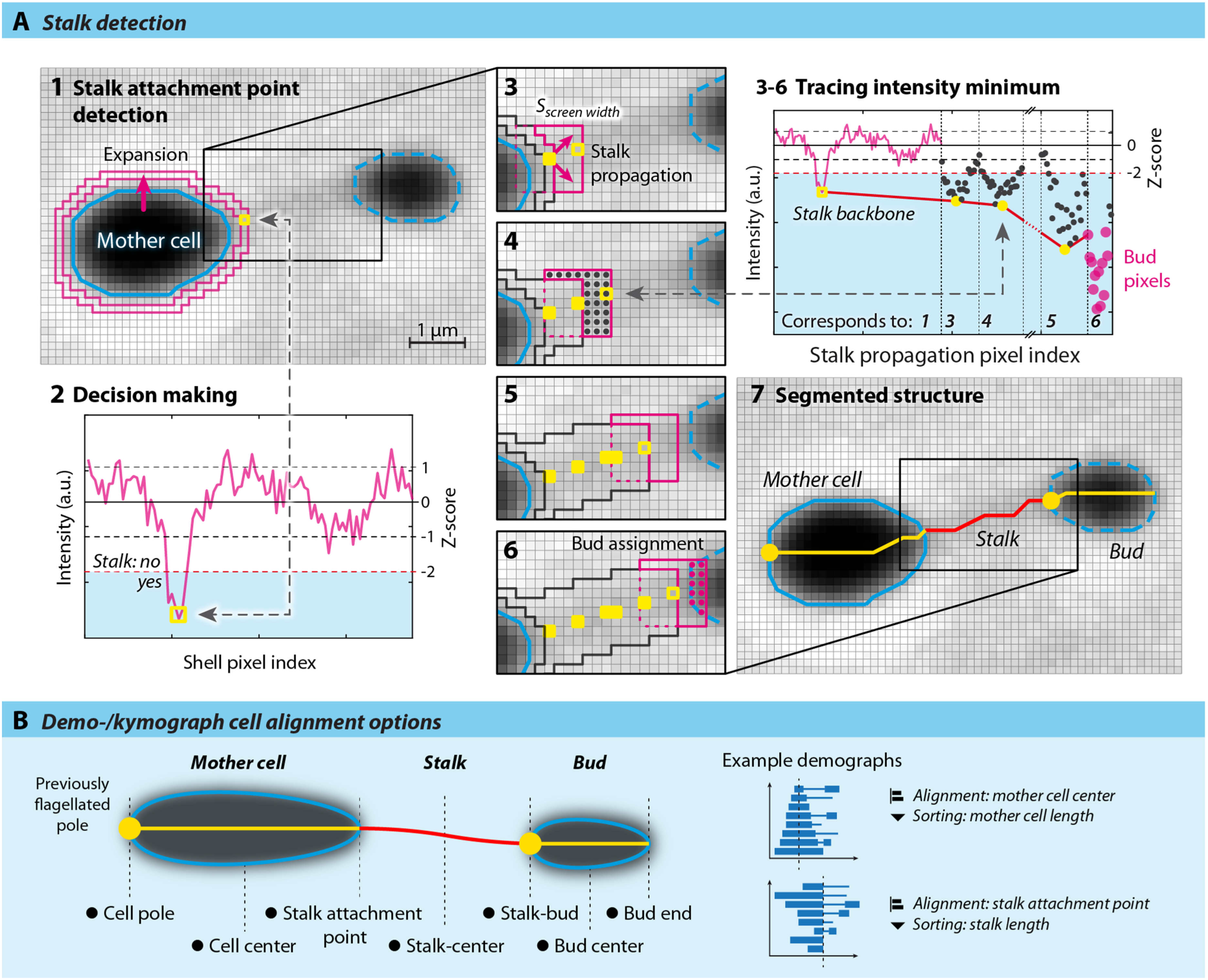
BacStaLk detects stalks in phase contrast images by tracing their subtle intensity patterns. It treats mother cells, stalks, and buds as connected morphological entities that can be appropriately aligned and sorted during quantitative data visualization. (A) Stalk detection algorithm. Steps 1-2: The stalk attachment point is determined by expanding each cell and identifying the pixel of lowest intensity (yellow pixel) inside the expanded cell-shell (magenta), whose intensity is below a typical z-score (number of standard deviations away from the mean). Steps 3-6: The stalk backbone is constructed by dilating the stalk attachment point by the stalk screening width and finding the next pixel of intensity below a typical z-score in the newly expanded area (magenta). The dilation is repeated for each valid backbone pixel and stopped if the stalk becomes too faint or another cell is encountered. Step 7: If the encountered cell is smaller than the initial cell, it is classified as a bud to the larger mother cell. If it is larger, the relation is inverted. (B) For visualization in demo-/kymographs, the cells can be aligned to all relevant positions (marked with a solid black dot) and sorted according to any of the extracted parameters.

We tested BacStalk on several species of stalked bacteria to verify the robustness of the stalk detection algorithm. The algorithms perform well for *C. crescentus* (Fig. 2A), *Brevundimonas aveniformis* (Fig. 2B), and *H. neptunium* (Figs. 3-4). To investigate whether stalks of different lengths are reliably detected, we imaged *C. crescentus* cells grown in the presence or absence of phosphate, as it has previously been shown that *C. crescentus* cells strongly elongate their stalk when starved for phosphate (34). Analysis of 500 cells grown in these two conditions showed that stalks of cells grown with phosphate were indeed considerably shorter and less variable in length (1.47 ± 0.75 μm) than stalks of cells deprived of phosphate (6.45 ± 2.90 μm) (Fig. 2A). Interestingly, we found that in the latter condition the cell length correlates at least partially with the stalk length. Stalk detection also works in extremely low contrast conditions in which the stalk is hard to distinguish from the background by eye, as generally observed for *B. aveniformis* (Fig. 2B). Here, 62% of all cells are stalked (n = 150), which is comparable to the 46% described in a previous study (35). Furthermore, our analysis showed that cells with stalks display a slightly larger average length of the cell body (2.47 ± 0.56 μm) than cells without stalks (2.05 ± 0.45 μm).

**FIG 2.**
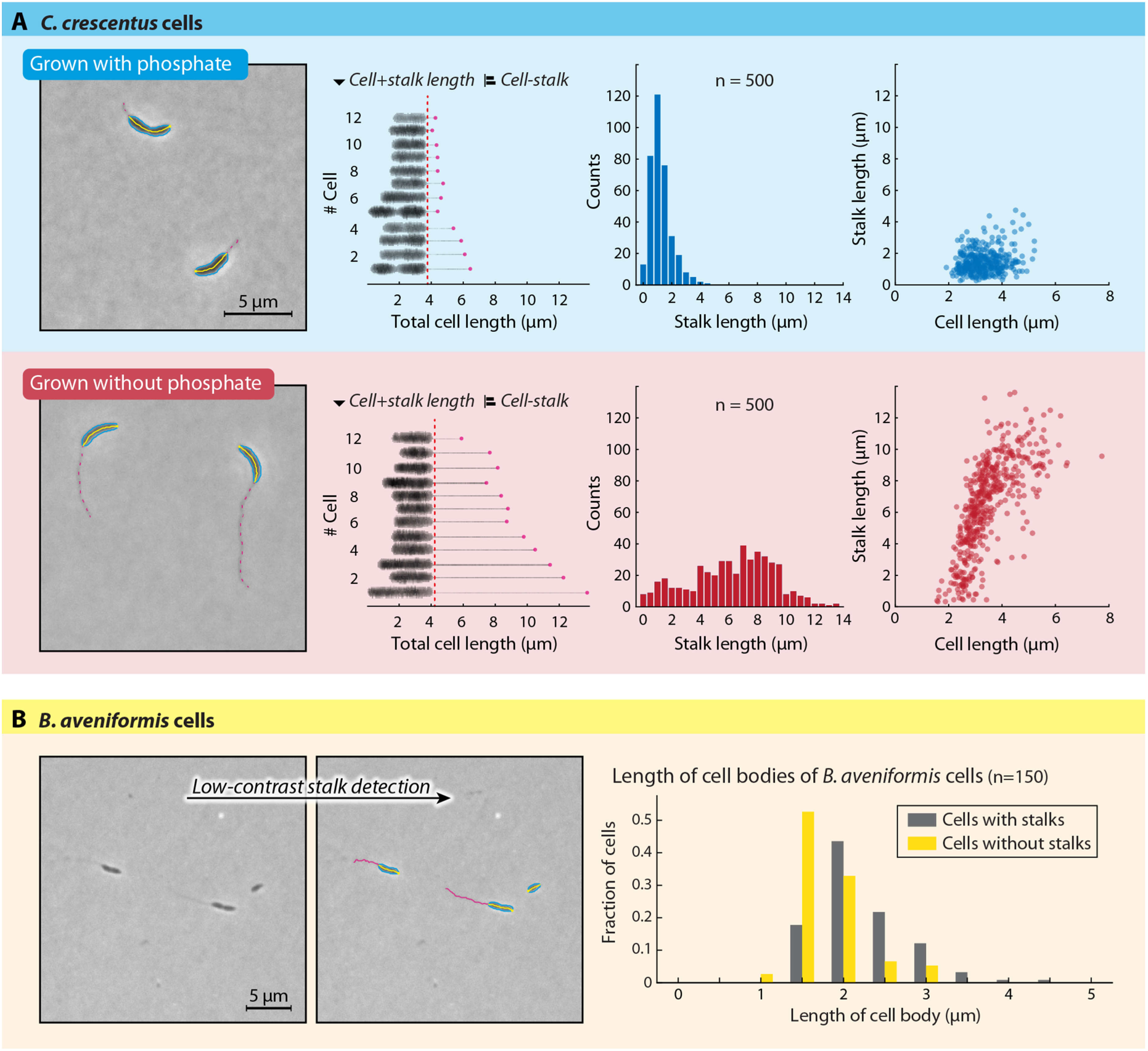
BacStaLk detects cell bodies, stalks, and buds in multiple species of stalked bacteria. (A) *C. crescentus* cells with normal and elongated stalks are detected and characterized separately. Screenshots created by BacStalk show cell outlines in light blue, medial axes in yellow, and stalks as magenta lines. Two-dimensional demographs display 12 representative cells for both conditions, aligned at the stalk attachment point and sorted according to the cell+stalk length after straightening the cells by medial axis coordinate system transformations. The distributions of stalk lengths are displayed in histograms. In addition, the correlations between the lengths of the cell bodies and the corresponding stalks are visualized *via* scatterplots. (B) The stalk detection algorithm also recognizes stalks that are hardly visible by eye, as demonstrated on *B. aveniformis* cells. Screenshots created by BacStalk are as described in panel A. Cell lengths of cells with and without stalks are displayed in a histogram.

**FIG 3.**
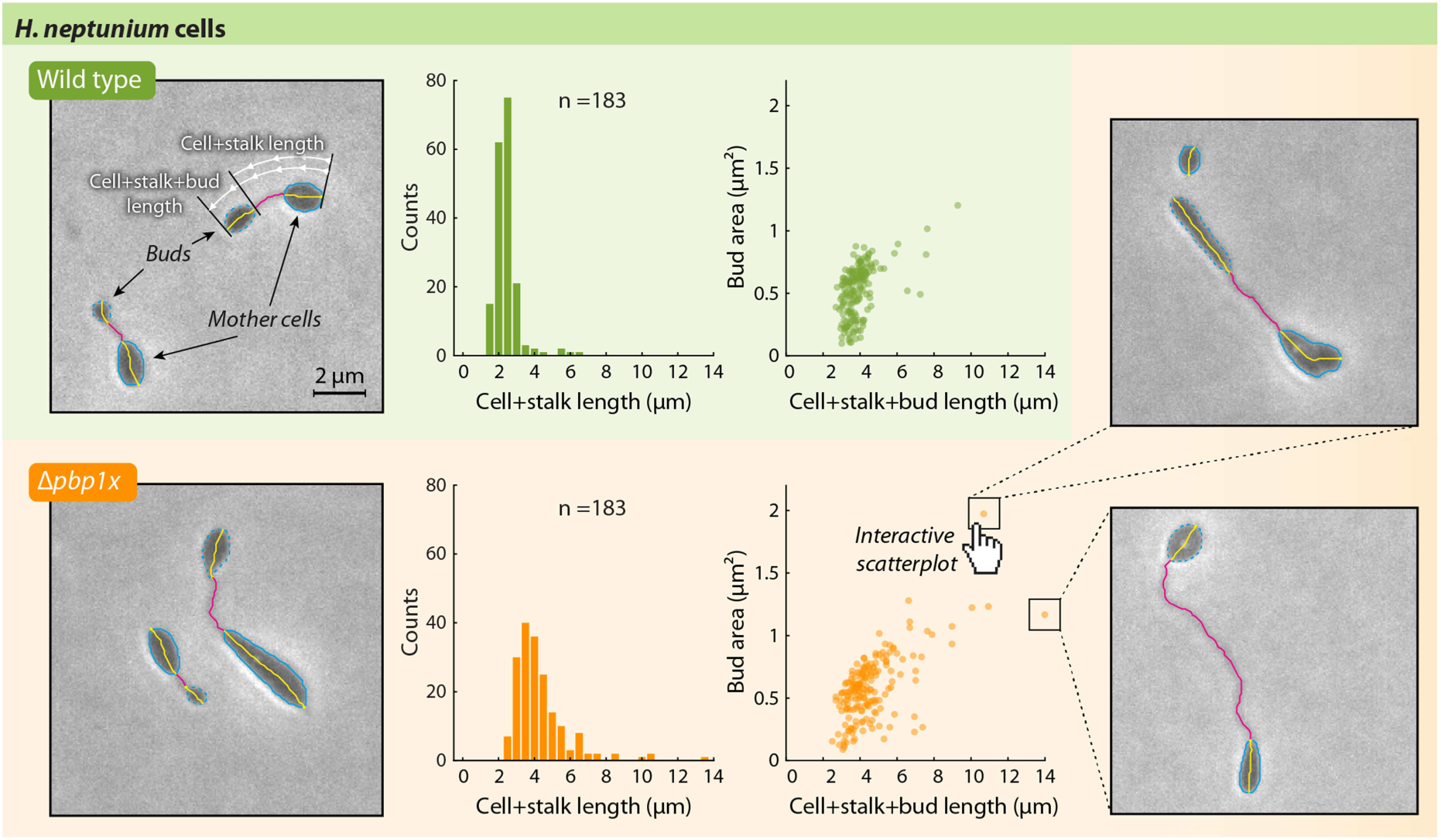
BacStaLk can be used for the analysis of mutants that differ in cell shape. Morphological differences between *H. neptunium* wild type and Δ*pbplx* cells are detected by quantitative analysis. In screen shots created by BacStalk, cell outlines are indicated by light blue lines, with bud cells highlighted by dashed outlines and medial axes in yellow. Upon deletion of *pbplx*, the cell+stalk length increases and shows a broader distribution, as shown in histograms. In addition, the population contains more cells with a larger bud area, which correlates with an increased total cell (cell+stalk+bud) length, as demonstrated by scatterplots comparing the bud area to the total cell length. The scatter plots generated by BacStalk are interactive; the underlying cell image is displayed and highlighted in the user interface, when clicking on a particular data point, e.g. in the scatter plot.

**FIG 4.**
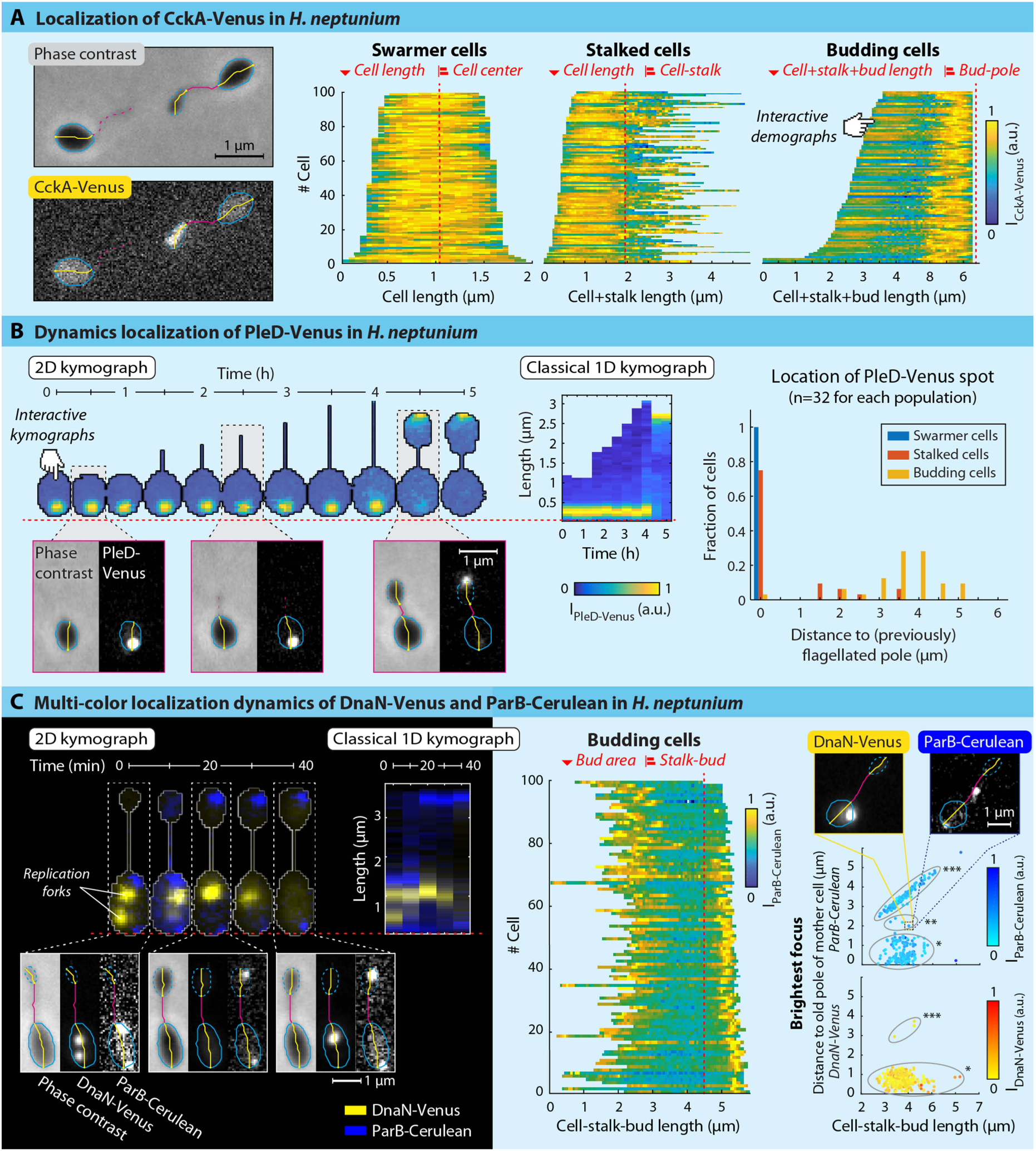
BacStaLk enables comprehensive and intuitive visualization of complex fluorescence patterns. (A) Analysis of the intracellular Localization of CckA-Venus in *H. neptunium* cells at different stages of development. Cells are aligned and ordered according to biologically relevant parameters (see red text above each demograph). The fluorescence profiles represent the mean fluorescence values along the medial axis after background subtraction and normalization such that the maximum fluorescence of each cell is equal. (B) Dynamic localization of PleD-Venus in *H. neptunium*, displayed as a 2D-kymograph (left) and a classical 1D-kymograph (middle) after background subtraction and normalization of the maximum fluorescence signal per time point. The histogram (right) shows the distance of the pixel with maximum PleD-Venus intensity from the old pole of the mother cell for swarmer cells, stalked cells, and budding cells. (C) The relative dynamics of multiple fluorescence signals in *H. neptunium* are visualized by 2D and 1D (classical) kymographs. The 2D kymograph clearly reveals DNA transfer to the bud (illustrated by the movement of blue ParB-Cerulean signal) before the entire DNA replication process is completed (shown by delocalization of the yellow DnaN-Venus signal). The fluorescence of ParB-Cerulean was corrected for the background and normalized (as the Cerulean signal bleached quickly), whereas the signal of DnaN-Venus was not background-corrected (in order to illustrate the difference between association and dissociation of DnaN-Venus from the chromosome). Demograph: Cells are aligned by the stalk-bud connection point and sorted according to the bud area (see red text above the demograph), revealing that ParB-Cerulean localization to the bud requires a minimum bud area. Scatterplots: ParB-Cerulean is present either in the mother cell (^⋆^) or at the flagellated pole of the bud (^⋆⋆⋆^) and only in rare cases in the stalk (^⋆⋆^). In contrast, the brightest DnaN-Venus focus is almost only present in mother cells (^⋆^). Dots in the scatterplot are colored relative to the fluorescence intensity of the brightest spot, showing that in the rare cases where the brightest DnaN-Venus focus is present in the bud (^⋆⋆⋆^), the DnaN-Venus signal was low (and thus replication was not ongoing).

In the stalked budding bacterium *H. neptunium*, BacStalk can accurately distinguish between mother cells, stalks, and buds for the wild type as well as for mutants with altered morphology (Fig. 3). Thus, BacStalk quantifies phenotypes in a more accurate, detailed, and faster manner than the current standard in the field (i.e. quantification *via* manual measurements). The power of such automated quantifications is illustrated by the fact that the cell morphology phenotype of the *H. neptunium* Δ*pbp1x* mutant (*i.e.* an increased length of budding cells), which was described qualitatively before (Cserti et al., 2017), can now be precisely quantified (Fig. 3): the combined length of mother cells and stalks was 3.2 ± 1.3 μm long for Δ*pbp1x* cells and 2.7 ± 0.7 μm for wild-type cells (both n = 183). The quantitative analysis by BacStalk also provided additional new insights. For instance, BacStalk revealed that the distribution of stalk lengths is dramatically broadened in the Δ*pbplx* background, and that bud area correlates partially with the total length of the concatenated entity of mother cell, stalk, and bud.

The combination of features offered by BacStalk facilitates detailed analyses of protein localization experiments based on fluorescence signals. To visualize protein localization, BacStalk can display combined intensity profiles of the mother cell, stalk, and bud, which can be aligned to according to relevant cellular landmark locations and sorted by any measured property of the combined structure (Fig. 1B). For generating intensity profiles along the medial axis of a cell, BacStalk fits a mesh of evenly spaced lines into each cell perpendicular to its medial axis, similar to Oufti (28). For each point on the medial axis, the mean or maximum of the intensity values along the corresponding line is calculated to obtain smooth intensity profiles along the cell medial axis. In addition, the mesh is used to perform a medial axis coordinate system transformation in order to reorient curved cells such that they can be arranged next to each other in two-dimensional (2D) demo- and kymo-graphs to conserve the full spatial information of patterns inside the cells (Fig. 2A, Fig. 4B,C, Fig. 5B). These 2D demo- and kymographs intuitively visualize the imaging data and provide important additional information in cases where the fluorescent protein of interest is located away from the medial axis. Furthermore, the intensity profiles can be normalized and background-corrected per cellular entity.

**FIG 5.**
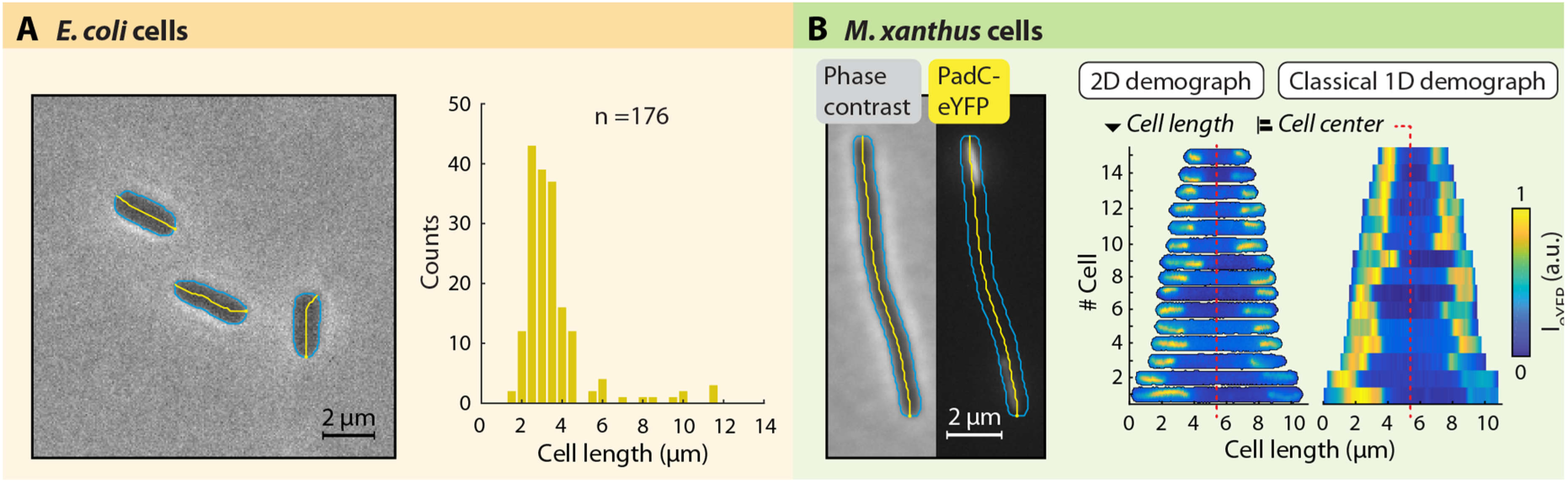
BacStalk is also a powerful tool for the analysis of non-stalked bacteria. (A) Images and histograms showing cell Lengths of *E. coli* TOP10 cells. (B) Images and 1D/2D demographs of *M. xanthus* cells (strain MO039), demonstrating the localization of PadC-eYFP to the subpolar regions.

BacStalk provides a very flexible visualization environment for protein localization experiments. The 1D and 2D demo- and kymographs, the use of concatenated intensity profiles, the possibility to align these profiles based on morphologically relevant criteria (e.g. alignment at the junction of the mother cell and stalk), and the option to display specific subsets of cells (i.e. swarmer cells, stalked cells without buds, and budding cells) are instrumental in understanding the localization behavior of proteins in different cell types in a mixed population. This is exemplified by an analysis of the localization dynamics of the histidine kinase CckA in *H. neptunium* using BacStalk, which verified the cell cycle-dependent localization previously observed in a qualitative manner by Leicht *et al.* (in preparation). BacStalk provides a means to automatically plot the intracellular localization of CckA-Venus in demographs separately for swarmer cells, stalked cells without buds, and budding cells (Fig. 4A). All demographs were sorted according to the length of the cellular structures: the first demograph is sorted by the length of the cell, the second by the combined lengths of mother cell and stalk, and the third by the combined lengths of mother cell, stalk, and bud. Cells were aligned at the cell center, the cell-stalk junction, or the bud pole opposite to the stalk, respectively.

Apart from its powerful visualization tools, BacStalk includes special features to track individual cells in time-lapse experiments and to analyze dynamic protein localization during stalk-terminal budding. As an example, we reinvestigated the cell cycle-dependent localization of the guanylate cyclase PleD in *H. neptunium* cells (Fig. 4B). The analysis by BacStalk provides a detailed quantification of the cell type-specific localization pattern that has previously only been described in a qualitative manner (36). In swarmer cells, PleD-Venus localizes at the flagellated pole of the mother cell. In stalked cells, by contrast, the PleD-Venus focus tends to be located at the tip of the stalk, whereas in most budding cells, it is detected in the bud at the pole opposite the stalk. Although the relocalization of PleD from the old pole of the mother cell to the old pole of the daughter cell can be traced in a 1D kymograph, the 2D kymograph additionally facilitates the correlation of protein translocation with cell morphogenesis (e.g. bud formation). Quantification of the localization of PleD-Venus in different cell types is also possible by determining and plotting its distance from the old pole of the mother cell (Fig. 4B).

To investigate the patterns of multiple fluorescence signals at different wavelengths simultaneously, BacStalk can create multi-channel kymographs (or demographs): the kymograph in Fig. 4C shows *H. neptunium* cells undergoing replication, in which (i) the replisome component DnaN is tagged with the fluorescent protein Venus and (ii) the origin of replication is followed with the help of a ParB-Cerulean fusion, which binds *parS* sites near the chromosomal origin of replication (Jung et al., in preparation). This two-color approach reveals the relative timing of replisome movement and origin segregation, and demonstrates that the origin of replication already moves to the bud before replication is completed (as visualized by delocalization of DnaN-Venus). In addition, the 2D kymograph visualization clearly identifies both replication forks as separate entities (DnaN foci inside first cell in kymograph, Fig. 4C). As in Fig. 4B, we determined the distance of the ParB-Cerulean and DnaN-Venus foci to the old pole of the mother cell. Using this representation, Fig. 4C shows that the replication origin (tagged by ParB-Cerulean) is only transferred to the bud once a certain bud size has been reached, and that the process of origin movement through the stalk must be fast, as the origin was captured inside the stalk in only ~1% of all analyzed cells (4/378).

BacStalk provides several analysis tools that greatly simplify data exploration and visualization. Similar to MicrobeJ (29), all plots created with BacStalk are interactive: clicking on a data point in a scatter plot (Figs. 2-4) or on a fluorescence profile in a demo- or kymograph (Figs. 4-5) displays the underlying cell, so that its raw image data and phenotype can be assessed. Furthermore, the output images of BacStalk showing the results of analyses or images of cells are publication-ready: all images in Figs. 2-5 have only been minimally edited after their export from BacStalk (e.g. change of background color, cropping of images, minor editing of the axes).

The features of BacStalk exemplified above for stalked bacteria, most notably its interactivity, its ease of use (see Fig. 6 for a description of the workflow), the one-click generation of 1D and 2D kymo- and demographs, and the ability to output publication-ready images are also applicable to the investigation of classical, non-stalked model organisms such as *E. coli* and *M. xanthus* (Fig. 5). The generation of interactive 2D demographs is a feature that is not available in any other image analysis software package. Its usefulness is demonstrated by an analysis of the localization dynamics of YFP-tagged PadC, an adapter protein connecting the chromosome partitioning ATPase ParA to subpolarly located bactofilin polymers in *M. xanthus* (37). Here, the 2D representation provides important information about the spatial arrangement of the filaments in the cell that cannot be appreciated in the standard 1D demo-graphs (Fig. 5B). As BacStalk is designed for micrographs with moderate to low cell densities, in which cells can be analyzed as isolated individuals, the software automatically excludes clumps of cells, because no cell-splitting functionality is implemented for the analysis of dense communities.

**FIG 6.**
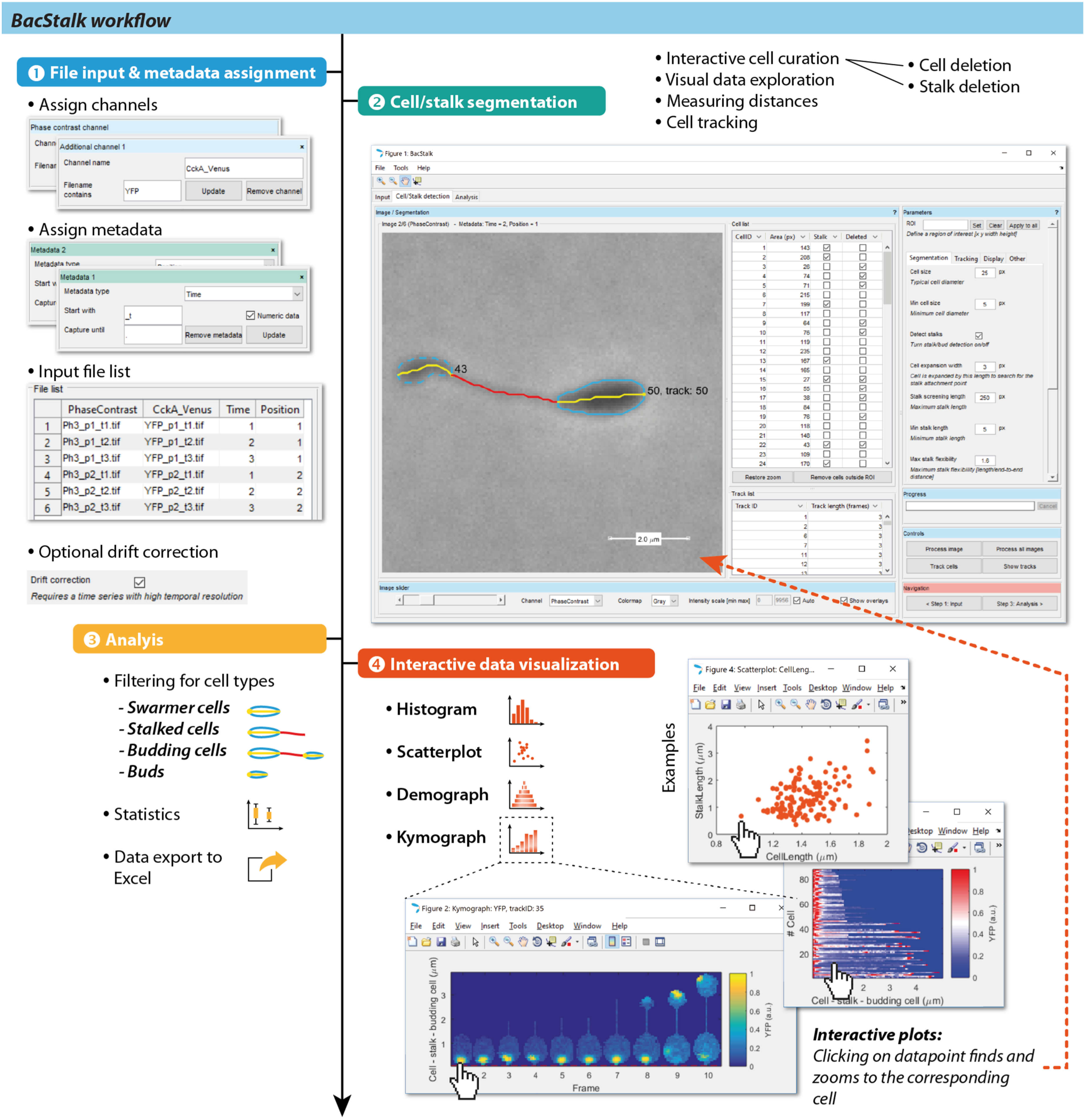
Typical workflow in BacStaLk. Step 1: Channel and metadata information are extracted from filenames based on keywords. For time-resolved image series, sample drift correction (image registration) can be performed. Step 2: BacStalk provides a powerful customizable image viewer for visual data exploration, cell and stalk segmentation, curation, and tracking. Image processing is optionally sped up by using MatLab’s Parallel Computing Toolbox. Step 3: The analysis tab provides tools for creating subsets of the bacterial population, calculating statistical parameters, and exporting the data to separate spreadsheet applications such as Microsoft Excel. Step 4: With a single click, complex data visualizations can be created, which are interactive and easily customizable using MatLab’s built-in plot editor to generate publication-ready vector graphics figures.

BacStalk was written in MatLab to make use of its built-in figure customization capabilities for generating publication-ready vector graphics and to provide advanced users with easy access to the underlying processed data. However, our main goal in the design of the software was to make it as user-friendly as possible and applicable on first-try without any programming knowledge. This ease of use is achieved by BacStalk’s powerful and fast graphical user interface (Fig. 6). In addition, the user is supported by a comprehensive documentation and detailed video tutorials, which are available online at http://drescherlab.org/bacstalk together with the open source code and stand-alone pre-compiled versions for various operating systems (Windows, Mac, and Linux). Overall, BacStalk facilitates high-throughput, in-depth, single-cell image analysis of stalked and non-stalked bacteria. It thus enables the study of many interesting stalked bacteria as novel model organisms, provides tools for more detailed analyses of established model organisms, and therefore constitutes an indispensable tool for bacterial cell biology.

## MATERIALS AND METHODS

### Bacterial strains

All strains analyzed in this study are listed in Suppl. Table 1. The plasmids and oligonucleotides used for their construction are listed in Suppl. Tables 2 and 3.

### Cultivation of cells

*C. crescentus* cells were grown in peptone-yeast-extract (PYE) medium (20), subsequently diluted 1/20 and grown for 24 h in M2-glucose (M2G) minimal medium (38) medium with phosphate or without phosphate (M2G^-P^) to induce stalk elongation. *B. aveniformis* cells were grown in PYE medium and *H. neptunium* cells in MB medium (Difco Marine Broth 2216, BD Biosciences, USA). *B. aveniformis*, *C. crescentus*, and *H. neptunium* cells were grown at 28 °C while shaking at 210 rpm. *E. coli* cells were grown in LB medium containing 30 μg/ml kanamycin at 37 °C while shaking at 210 rpm. *M. xanthus* cells were grown in CTT medium (39) containing 50 μg/ml kanamycin at 32 °C while shaking at 210 rpm. The synthesis of PadC-eYFP was induced for 3 h with 5 μM vanillate. For imaging, cells were grown to early exponential phase (OD_600_ = 0.2-0.4) and spotted on pads consisting of 1% agarose (peqGOLD Universal Agarose, peqlab, Germany) and the respective medium. In the case of *H. neptunium* strain JR47, the cells were imaged on pads prepared with 4-fold diluted MB medium, whereas *M. xanthus* MO039 cells were applied to pads made with 5-fold diluted CTT, in both cases to decrease background fluorescence.

### Light microscopy

All strains, except for *E. coli* and the *H. neptunium* wild-type and Δ*pbp1x* strains, were analyzed with a Zeiss Axio Observer. Z1 inverted microscope (Zeiss, Germany) equipped with a Zeiss Plan-Apochromat 100x/1.4 Oil Ph3 objective, Chroma ET-YFP and ET-CFP filter sets, and a pco.edge 4.2 mHQ camera (PCO, Germany). *E. coli* and *H. neptunium* wild type and Δ*pbp1X* cells were imaged with a Nikon Ti-E inverted epifluorescence microscope equipped with a Nikon Plan-Apochromat λ100x/1.45 Oil Ph3 objective and an Andor Zyla 4.2plus sCMOS camera.

## ACKNOWLEDGEMENTS

We are grateful to Oliver Leicht, Emöke Cserti, Alexandra Jung, Lin Lin, Manuel Osorio Valeriano, and Julia Rosum for the construction of plasmids and strains used in this study. Revathi Lakshmi Pulpetta is acknowledged for imaging strain JR47 and Manuel Osorio Valeriano for imaging *M. xantus* cells. In addition, we would like to thank Adrian Izquierdo Martinez, Manuel Osorio Valeriano, Revathi Lakshmi Pulpetta and Till, as well as the attendants of the BacStalk workshop in Marburg for their feedback after testing BacStalk prior to release.

---

**SUPPL. TABLE 1.**
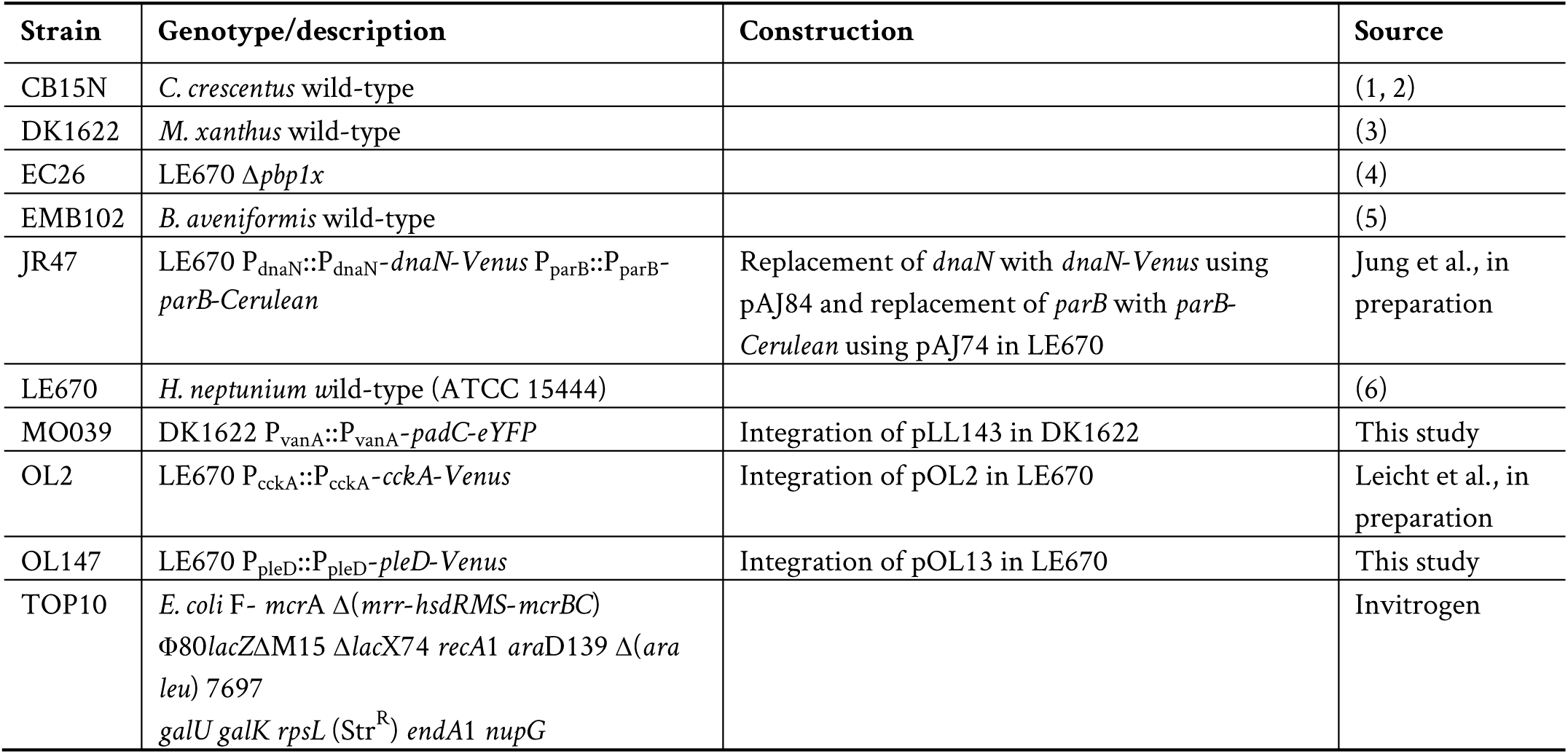
Strains used in this study.

**SUPPL. TABLE 2.** Plasmids used in this study.

| Plasmid | Description | Construction | Source |
| --- | --- | --- | --- |
| pAJ74 | pNPTS138 derivative for generating an in-frame replacement of *parB<sub>HN</sub>* with *parB<sub>HN</sub>*-Cerulean | a) PCR amplification of the last 965 bp of *parB<sub>HN</sub>* excluding stop-codon fused to Cerulean with primers oAJ299 and oAJ257<br>b) PCR amplification of 838 bp downstream of *parB<sub>HN</sub>* using oAJ300 and oAJ301<br>c) Digestion of the products with HindIII, KpnI and NheI<br>d) Ligation of the fragment into HindIII/NheI-treated pNPTS138 | Jung et al., in preparation |
| pAJ84 | pNPTS138 derivative for generating an in-frame replacement of *dnaN<sub>HN</sub>* with *dnaN<sub>HN</sub>*-Venus | a) PCR amplification of the last 1116 bp of *dnaN<sub>HN</sub>* excluding stop-codon fused to Venus with primers oAJ345 and oAJ315<br>b) PCR amplification of 927 bp downstream of *dnaN<sub>HN</sub>* using oAJ343 and oAJ344<br>c) Digestion of the products with HindIII, NdeI and EcoRI<br>d) Ligation of the fragment into HindIII/EcoRI-treated pNPTS138 | Jung et al., in preparation |
| pLL143 | pMR3690 bearing *padC*-eYFP<sub>MX</sub> |  | (7) |
| pMR3690 | Vector for the expression of a gene under the control of the *vanA* promoter, integrating at the *M. xanthus* Mxan18_19 chromosomal locus, Kan<sup>R</sup> |  | (8) |
| pNPTS138 | *sacB*-containing suicide vector used for double homologous recombination, Kan<sup>R</sup> |  | M. R. Alley, unpublished |
| pOL2 | pVENC-2 bearing *cckA<sub>HN</sub>* | a) PCR amplification of the last 613 bp of *cckA<sub>HN</sub>* excluding stop-codon with primers oJK177 and oJK178<br>b) Digestion of the product with NdeI and EcoRI<br>c) Ligation of the fragment into NdeI/EcoRI-treated pVENC-2 | Leicht *et al.*, in preparation |
| pOL13 | pVENC-2 bearing *pleD<sub>HN</sub>* | a) PCR amplification of the last 609 bp of *cckA<sub>HN</sub>* excluding stop-codon with primers oOL15 and oOL16<br>b) Digestion of the product with NdeI and KpnI<br>c) Ligation of the fragment into NdeI/KpnI-treated pVENC-2 | This study |
| pVENC-2 | Integration plasmid to fuse 3' end of a target gene to *venus*, Kan<sup>R</sup> |  | (9) |

**SUPPL. TABLE 3.**
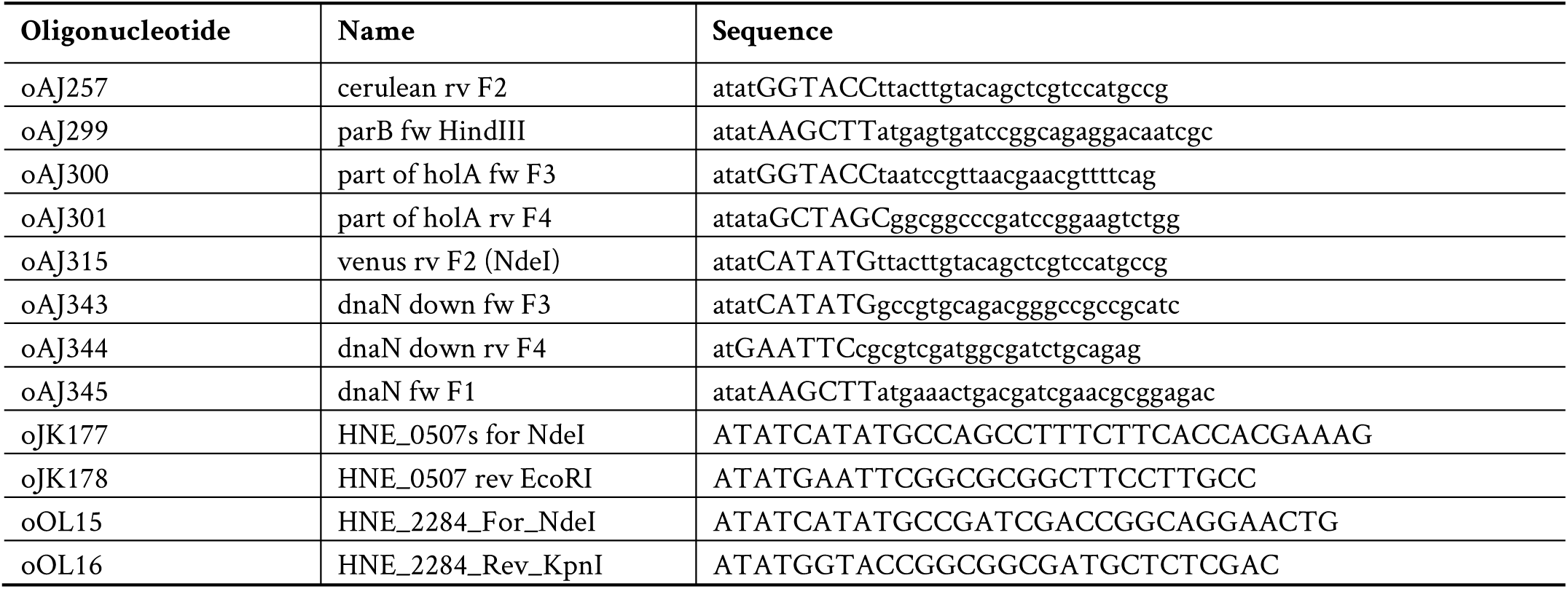
Oligonucleotides used in this study.

